# Three-Dimensional (3D) Fibronectin Nano-Array Presented on Fibrin Matrix Accelerates Mice Skin Wound Healing

**DOI:** 10.1101/2020.05.04.077891

**Authors:** Carlos Poblete Jara, Ou Wang, Thais Paulino do Prado, Ayman Ismail, Frank Marco Fabian, Han Li, Licio A. Velloso, Mark A. Carlson, William Burgess, Yuguo Lei, William H. Velander, Eliana P. Araújo

## Abstract

Plasma fibrinogen (F1) and fibronectin (pFN) polymerize to form a fibrin clot that is both a hemostatic and provisional matrix for wound healing. About 90% of plasma F1 has a homodimeric pair of γ chains (γγF1) and 10% has a heterodimeric pair of γ and more acidic γ’ chains (γγ’F1). We have synthesized a novel fibrin matrix exclusively from a 1:1 (molar ratio) complex of γγ’F1 and pFN in the presence of highly active thrombin and recombinant Factor XIII (rFXIIIa). In this matrix, the fibrin nanofibers were wrapped with periodic 200-300 nm wide pFN nanobands (termed γγ’F1:pFN fibrin). In contrast, fibrin made from 1:1 mixture of γγF1 and pFN formed a sporadic distribution of “pFN droplets” (termed γγF1 +pFN fibrin). The γγ’F1:pFN fibrin enhanced the adhesion of primary human umbilical vein endothelium cells (HUVECs) relative to the γγF1+FN fibrin. Three dimensional (3D) culturing showed that the γγ’F1:pFN complex fibrin matrix enhanced the proliferation of both HUVECs and primary human fibroblasts. HUVECs in the 3D γγ’F1:pFN fibrin exhibited a starkly enhanced vascular morphogenesis while an apoptotic growth profile was observed in the γγF1 +pFN fibrin. Relative to γγF1 +pFN fibrin, mouse dermal wounds that were sealed by γγ’F1:pFN fibrin exhibited accelerated and enhanced healing. This study suggests that a 3D pFN nano-array presented on a fibrin matrix can promote wound healing.

## Introduction

Skin wound healing consists of four phases: hemostasis, inflammation, proliferation, and remodeling. After an injury in healthy people, plasma fibrinogen (F1) and plasma fibronectin (pFN) (10:1 at molar ratio) are polymerized by thrombin and factor XIII (FXIII) to form a fibrin clot to stop bleeding (i.e. the hemostasis)^1^. In the clot, fibronectins are attached covalently and uniformly to the fibrin nanofibers^2^. In a material sense, F1 and pFN are both diverse macromolecules that greatly contrast each other in structure. F1 is a linear, hexameric, 340 kDa glycoprotein having 2 alpha, 2 beta, and 2 gamma chains^3–5^ pFN is a 440 kDa globular protein having two chains with cellular binding domains. Both F1 and pFN have cellular binding domains that remain cryptic until crosslinking by activated Factor XIII (FXIIIa) into the fibrin matrix^6^. About 90% of F1 circulates as a homodimeric pairing of γ-chains (γγF1) while the remaining 10% presents a heterodimeric pairing of γ with a more acidic γ’-chain (γγ’F1)^6^. The γ’-chain occurs from an alternative transcript having 20 additional amino acids at the carboxy-terminal that has been correlated with a pleiotropic impact on vascular health^6–8^. Importantly, pFN and γγ’F1 both occur at about the same concentration in plasma of about 300 μg/mL^7–11^ and therefore their presence in fibrin clots.

The hemostasis begins with thrombin-mediated activation of F1 and FXIII. The activated F1 produces a semi-soluble, viscoelastic fibrin aggregate^3,12,13^ containing pFN entrained from plasma. The FXIII is inherently present within the polymerizing fibrin aggregate as a complex with 1 out of every 100 γγ’F1 molecules^7^. It disassociates upon the formation of activated FXIII that enables efficient intra-fibrin cross-linking activity to produce an insoluble fibrin matrix^14–18^, and to simultaneously anchor the matrix to the wound surface in about 3 minutes^7,19^. This clotting process causes protein conformational changes that induce interactions between pFN, fibrin, and also wound healing cells.

After hemostasis, the fibrin clot acts as a provisional matrix for different cell types to heal the wound^20^. Neutrophils and inflammatory (or M1) macrophages first enter the matrix to clear bacteria and tissue debris. Neutrophils secrete anti-microbial peptides, proteases and reactive oxygen species (ROS) to remove bacteria and damaged tissue^21^. These chemicals are toxic to surrounding healthy cells as well^21^. The M1 macrophages phagocytosis bacterial and damaged tissue^22–30^. Both cell types directly bind to fibrin nanofibers via their αMβ2 integrin receptors. Once the wound is debrided (takes ~3 days^31^), neutrophils undergo apoptosis and M1 macrophages switch to the regenerative M2 phenotype that secretes cytokines and growth factors to recruit and stimulate healing cells such as endothelial cells, fibroblasts and keratinocytes to heal the wound (i.e. the proliferation phase)^32–34^. Endothelial cells form new blood vessels and fibroblasts deposit new extracellular matrix proteins to form a granulation tissue (takes ~7 days)^32–34^. Keratinocytes migrate on the granulation tissue to form a new epidermis to close the wound (i.e. re-epithelization)^33^. Healing cells adhere to the fibrin matrix by binding to fibronectins with their integrin receptors^32–35^. The binding also triggers downstream signalings that are required for these cells to migrate, proliferate, and repair the wound^32–36^. Fibronectins also bind, enrich, protect, orientate, and present growth factors locally to potentiate their functions^37–48^. In short, the fibrin-based provisional matrix is the central stage for cells to heal a wound, and fibronectins are essential for adhering and stimulating the healing cells. Consequently, insufficient provisional matrix or fibronectins or fibronectin-induced cell signalings lead to pathological wound healings^32–35,37^. For instance, small wounds can heal in about 14 days, while large wounds take much longer time to heal partially due to the insufficient fibrin-fibronectin provisional matrix.

We hypothesize that increasing the pFN content or FN-cell interaction in the fibrin matrix may promote wound healing. As said, the natural fibrin matrix only contains 10% (molar ratio) fibronectins^1^. Previously, we reported a novel method to increase the pFN content up to 50% (molar ratio) in the fibrin matrix and to present the pFN as nanobands on the fibrin nanofibers^8^. In this report, we show that this novel pFN: γγ’F1 fibrin accelerates re-epithelization and granulation tissue formation in mouse dermal wounds. This engineered fibrin sealant has attributes useful to both hemostasis and to promote faster wound healing including improvement of granulation tissue quality.

## Methods

### Materials

All reagents were obtained from Sigma Chemical Company (St. Louis MO) unless otherwise noted. Human source plasma was provided by the U.S. Army Materials Command (Fort Detrick, MD.). Recombinant human thrombin was purchased from ZymoGenetics (Seattle, WA). DEAE Sepharose fast flow, Superose 6, and Gelatin Sepharose were purchased from GE Healthcare (Uppsala, Sweden). 4-12% NuPage Bis-Tris SDS polyacrylamide gels, Colloidal Blue stain and See Blue molecular weight markers were from Invitrogen (Carlsbad, CA). Polyclonal antibody (GMA-034) for human fibrinogen was purchased from Green Mountain Antibodies (Burlington, VT). Antibody (ab2413) for human fibronectin was from Santa Cruz Biotechnology.

### Production of recombinant Factor XIII

Recombinant active Factor XIII (rFXIIIa) was expressed in *Pichia Pastoris* following published methods^49–51^. The activity was determined using Pefakit (Pentapharm, Norwalk, CT).

### Preparation of fibrinogen and fibronectin from human plasma^8^

Three units of human plasma that had been stored at −80 °C were thawed at 4 °C. The plasma was centrifuged at 4000 rpm for 20 min. The cryoprecipitate was re-suspended in 45 mM sodium citrate, 100 mM 6-aminocaproic acid, pH 7.0 at 37 °C. The solution was then centrifuged for 25 min at 4000 rpm. The supernatant was treated with a solvent detergent to inactivate the virus by addition of 0.15% TnBP, 0.5% Triton X-100 and stirred at room temperature for 60 min. The supernatant was adjusted to 1 M ammonium sulfate. The sample was stirred at room temperature for 30 min and then centrifuged at 2000 rpm for 15 min at room temperature. The pellet was re-suspended in 20 mM sodium citrate, 100 mM NaCl. The sample was dialyzed overnight at room temperature against the same buffer. The dialyzed sample was centrifuged at 2000 rpm for 15 min at room temperature.

The supernatant was fractionated to isolate γγF1, γγ’F1 and pFN using DEAE Sepharose Fast Flow chromatography at room temperature^52^. The sample was applied to a DEAE column then washed with 10 column volumes of 20 mM sodium citrate, 100 mM NaCl, pH 7.4. The unabsorbed fraction was γγF1. Bound protein was eluted with a linear gradient of 0.1 M to 1.0 M NaCl. The pooled eluate produced an approximately equimolar mixture of γγ’F1 and pFN that was dialyzed against 20 mM citrate buffer, 20 mM NaCl at pH 6.8. The dialyzed mixture was concentrated using a 10 kDa centrifugal ultrafiltration device at 4000 g for 30 min.

Pure pFN and γγ’F1 were produced by application of the DEAE eluate pool to gelatin-Sepharose. Briefly, a 10 mL γγ’F1:pFN (“γγ’F1:pFN” that we will name from here as “complex”) was applied to a 3 mL analytical gelatin-Sepharose column. The column was washed with 10 volumes of the dialysis buffer then eluted with the same buffer containing 6 M urea. pFN was eluted by urea while the γγ’F1 fell through the column.

### 2D adhesion assay

Human foreskin fibroblasts were isolated and cultured in DMEM with 10% FBS. The use of primary human fibroblasts from anonymous donors was approved by the Institutional Review Board at the University of Nebraska Medical Center and by the Research and Development Committee at the Omaha VA Medical Center. Passage 4 to 6 cells were used. HUEVCs were purchased from Lonza and cultured in EGM2 medium. Passage 3 and 4 cells were used for the experiments. 100 μl fibrin matrix was made in one well of the 96-well plate. 1×10^4^ cells were placed in each well with 200 μl medium. 3 hours after plating, medium, and unattached cells were removed. Cells were washed with 200 μl PBS once. 100 μl fresh medium with 10 μl Alamar blue reagent was added and incubated for 3 hours. 100 μl medium was collected for measuring the fluorescence that was calibrated to a standard curve to calculate the numbers of attached cells within each well. After the quantification, cells were fixed with 4% PFA and stained with DAPI. The nuclei were recorded with a fluorescence microscope.

### 3D cell culture within the fibrin matrix

100 μl fibrin matrix with 2×10^4^ cells were made in individual wells of a 96-well format plate. 200 μl medium was added to each well. At varied time points after plating, cell morphologies were recorded by phase-contrast microscopy. The medium was removed and 100 μl fresh medium with 10 μl Alamar blue reagent were added and incubated for 3 hours. 100 μl medium was collected for-measuring the fluorescence that was calibrated to a standard curve to calculate the numbers of cells in each matrix. The viability of cells within the gel was accessed with a Live/Dead cell assay kit (Invitrogen) according to the Manufacturer’s instruction.

### Fibrin formulation

To make the fibrin matrix for the above cell culture experiment, the following components were mixed to the final concentration of fibrinogen (γγF1, γγ’F1) at 2.5 mg/mL; thrombin at 1 U/mL; rFXIII at 15.4 U/mL; and pFN at 3.3 mg/mL. F1 and pFN were at a 1 to 1 molar ratio. The mixtures were incubated at 37 °C for 15 min to form the fibrin matrix.

### SEM and confocal microscopy

Fibrin matrices were made according to the formulation above. For SEM, samples were fixed with 2.5% glutaraldehyde in 100 mM phosphate buffer (PH 7.0) at room temperature for 1 hour, then at 4 °C overnight. Samples were rinsed with phosphate buffer twice, 10 minutes each. Ethanol dehydration series were performed: 30%, 50%, 70%, 2 x 95%, 2 x 100%, 5 minutes for each procedure. Then samples were treated with hexamethyldisilazane (HMDS) as the following: 33% HMDS, 66% HMDS, 2 x 100% HMDS, 2 minutes for each procedure. Samples were left in 100% HMDS to air-dry at least overnight before sputter-coated with gold-palladium and imaged with the scanning electron microscope (Hitachi S4700 Field-Emission SEM, Hitachi, Tokyo, Japan) at 10 kV. For confocal microscopy, samples were fixed with 4% paraformaldehyde (PFA) at 4 °C overnight, washed with PBS for 3 times, and blocked with 5% goat serum for 1 hour before incubating with primary antibodies at 4 °C overnight. After extensive washing, secondary antibodies were added and incubated for 2 hours at room temperature. Samples were then washed with PBS before imaged with confocal microscopy (Nikon A1-R confocal system on a Nikon Eclipse 90i upright fluorescence microscopy).

### Animal research

All animal protocols were approved by the Institutional Animal Care and Use Committee of the State University of Campinas (protocol number 4467-1/2017) and the University of Nebraska-Lincoln (protocol 1571). All experimental procedures were performed following the guidelines of the Institutional Animal Care and Use Committee of the University of Campinas and the University of Nebraska-Lincoln and. For the surgical procedure, mice were anesthetized using intraperitoneal of xylazine (10 mg/kg) and ketamine (100 mg/kg). Individual doses were calculated using *Labinsane App* (Labsincel, 2017). A deep anesthetic plane was verified by the loss of pedal and corneal reflexes. Two symmetrical 6-mm full-thickness excisional wounds were created on the back of each mouse as described previously^53^. Donut-like splints with an 8 mm diameter of the center hole and 15-mm diameter of the disc were created. The splint was carefully placed around the wound and secured to the skin with eight interrupted sutures. After surgery, animals were maintained in a warm bed for anesthetic recovery. We applied intraperitoneal injection of tramadol (25 mg/kg) at 0, 12, and 24 h after surgery as an analgesic treatment.

### Mice fibrin sealant treatment

First, we cleaned the unwounded surrounding skin with a saline solution to remove any traces of biological residue. We sealed the wound with 50 μL fibrin. The fibrin sealant contained 2.5 mg/mL fibrinogen, 3.3 mg/mL fibronectin, 15.4U/mL rFXIIIa, 5 U/mL thrombin, 16 ng/mL VEGF, 5mM CaCl2 in the Ringer solution. Control animals received the same amount of CaCl2 and VEGF in the Ringer solution. The wounds were covered with a sterile transparent dressing. On days 2, 4, 6, and 9, we cleaned and disinfected the surrounding skin as described above and reapplied 25 μL fibrin sealant.

### Wound healing process documentation and analysis

Macroscopic evaluation of the healing process was evaluated by photographs taken at 0, 2, 4, 6, 9, and 11 days after wounding using a Nikon D610 (Nikon Systems, Inc., Tokyo, Japan) camera with an AF-S DX NIKKOR 18-55mm f/3.5-5.6G VR objective. The same distance (wound to the objective lens), top white fluorescence illumination, and operator were used on each occasion. We calculate Wound Area as follows: Wound Area (%) = [(wound area of day 0 - wound area of day x) / wound area of day 0] × 100. In each group, we use 16 to 20 photographs per day of analysis. The percentage of wounds with complete closure was calculated by the number of mice that achieve 100% of reepithelization over the total of mice. Wound re-epithelization was examined by ImageJ Software (1.49v).

### Histology

The skin samples containing wounded and unwounded areas were carefully harvested using an 8 mm biopsy punch. The samples were fixed in cold 4% (wt/vol) formaldehyde in PBS at 4 °C overnight. Formaldehyde was discarded and the tissues were washed 3 times using PBS 1x. We divided the circular sample in the middle of the circle to standardize the diameter of the sectioned samples. Then, the samples were treated with increasing concentration of alcohol (70%, 80%, 95%, and 100%) followed by xylol and paraffin and finally fixed in paraffin blocks. We sectioned at 5.0 μm and set 4-6 slices on microscope slides pretreated with poly-L-lysine. Following this procedure, we stained the skin sections with hematoxylin and eosin (Sigma). We incubated the slices with hematoxylin for 30 s, rinsed them in distilled water, incubated them for 30 s with eosin, washed them over in distilled water, and dehydrated them. For Picrosirius red staining (Picro Sirius Red Stain Kit, Sigma) we soaked the section for 30 min in distilled water. Then we incubated the section with Picro-Sirius red solution for 60 min. Slides were washed with acetic acid solution and then they were washed with absolute alcohol. For Masson’s Trichrome Staining we incubated slide in preheated Bouin’s Fluid for 60 min and cool for 10 min. We rinsed it in distilled water and then we incubated in Weigert’s Iron Hematoxylin for 5 min, Biebrich Scarlet Acid Fuchsin solution for 15 min, differentiate in the phosphomolybdic acid solution for 15 min, Aniline Blue solution for 10 min and acetic acid solution for 5 min. Finally, we mounted the slices on Entellan®, digitally captured the images under bright field microscopy (Leica DM4500B), and processed with Leica Application Suite 3v for *Full view reconstruction*.

### Collagen quantification

To analyze the birefringence patterns, we used microscopy (Nikon^®^ Corp., Kanagawa, Japan) with two CPL polarized filters. We captured the images with a 100x or 400x objective and assessed them for collagen Type III (bright green color) and collagen Type I (red color). The birefringence for Type I collagen was dense, strongly birefringent red fibers, whereas Type III collagen showed thin, green, and softly birefringents^54^. We used Image Pro Plus (version 4) to measure the stained collagen fibers as the percentage of total pixels in each wound pictures.

### Full view reconstruction

To increase the precision of evaluation, we reconstructed the whole skin section using unwounded and wounded images. We used a Nikon Microscopy to take 10-15 adjacent microphotographs (with 30% of common areas, 5x objective) and then, reconstructed each skin section. We applied the *Photomerge tool* with the *perspective* layout in Adobe Photoshop (19.1v).

### Transcript expression determination by real-time Polymerase Chain Reaction (PCR)

We used sample aliquots with 5.0 ng RNA for reverse transcription applying random hexamer primers and Superscript Moloney-MLV reverse transcriptase. PCR reactions were performed in real-time using the TaqMan^™^ system (Applied Biosystems). The GAPDH gene (TaqMan^™^—Applied Biosystems) was employed as the endogenous control for the reaction to which the expression levels of the gene of interest in different samples were normalized. After calculating the efficiencies of amplification, a scatter plot was constructed to define the series of concentrations for which the system was competent. Primers for the target genes were purchased from Applied Biosystems and IDT DNA Technology against the TGF-1β (Mm.PT.58.43.479940), TNF-α (Mm00443258_ m1), IL-6 (Mm00446190_m1), VEGF-α (Mm. PT.58.14200306), FGF-1 (Mm.PT.56a.41158563), IGF-1 (Mm.PT.58.5811533), IL-1β n(Mm00434228_m1), MCP-1 (Mm99999056_m1), IL-10 (Mm01288386_m1), SDF-1 (Mm00445553_m1), ITGA (NM_010577.3), Plaur (ENSG00000011422), MMP9 (ENSG00000100985), Procr (ENSG00000101000) FLRT2 (ENSG00000185070), F4/80 (Mm00802529_m1), Cd11b (Mm00434455_m1), iNos (Mm.PT.5843705194), Cd163 (Mm00474091_m1), Arg1 (Mm.PT.58.8651372) and GAPDH (4352339E). We Used the StepOnePlus Real-Time PCR System^™^ (Thermo Fisher Scientific). For transcript quantification, the PCR in realtime were carried out in duplicate in reactions containing 3 μl of TaqMan Universal PCR Master Mix 2X, 0.25 μl of primers and probe solution, 0.25 μl of water and 4.0 μl of cDNA. The negative control used 4.0 μl of water in place of cDNA. The cycling conditions were as follows: 45 cycles of 95 °C for 2 min, 95 °C for 5 s, and 60 °C for 30 s. The relative gene expression values were obtained by analyzing the results in the StepOne^™^ 2.2.2v (Applied Biosystems).

### Statistical analysis

The data are presented as the mean ± S.D. We used an unpaired t-test to compare two groups and one-way ANOVA to compare more than two groups. P < 0.05 was considered statistically significant. We used GraphPad Prism 6 for Windows 6.01v to perform statistical analysis.

## Results

### A bioprocess for purifying γγ’F1:pFN complex

Recently, we developed a scalable process to separate γγF1 and γγ’F1 from human plasma at production scale^8^. We observed that γγ’F1 and pFN isolated as a noncovalent complex at about 1:1 molar ratio^8^. This complex could be dissociated by passing through a gelatin affinity column where the pFN was absorbed and γγ’F1 passed through the affinity column.). We also found that the dissociated γγ’F1 and pFN could quickly re-associate at room temperature^55^. The high purity of the compositions used in this paper consisting of pFN, γγF1, γγ’F1, γγ’F1:pFN and a 1:1 mixture of γγF1 and pFN (γγF+pFN) were confirmed with reduced SDS-PAGE (**Fig. 1a**).

**Fig. 1.**
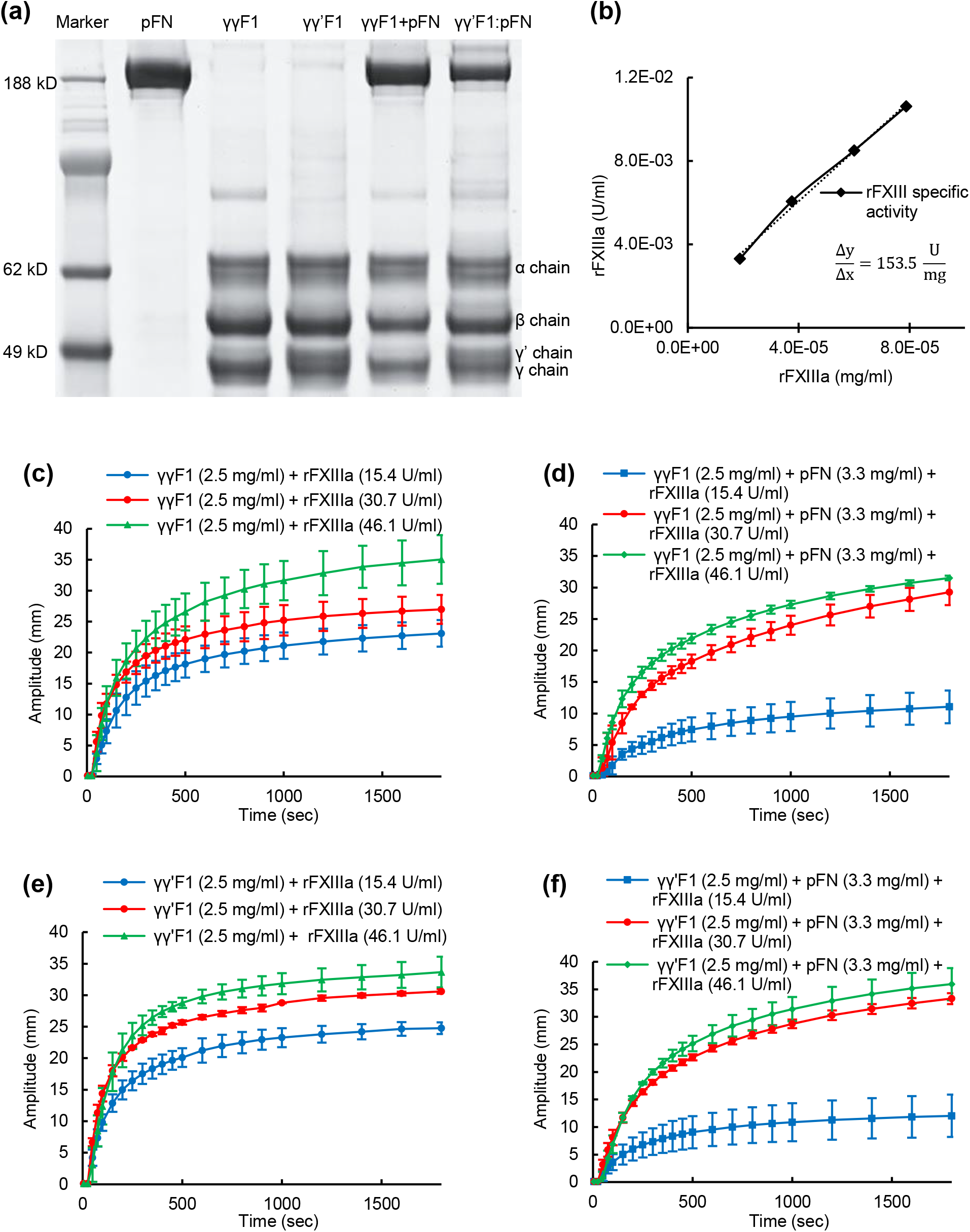
a) Reducing SDS-PAGE gel. b) rFXIIIa activity (153.5 ± 19 U/mg) was measured using the 5-(biotinamido) pentylamine incorporation assay. The effect of rFXIIIa on the viscoelastic and clotting kinetics of γγF1 (c), γγF1+pFN mixture (d), γγ’F1 (e), and γγ’F1:pFN complex (f) were evaluated via TEG assay.

### Highly active Factor XIII was critical to forming a stable fibrin matrix at low sealant concentration

The normal concentration of fibrinogen in blood plasma is 2-4 mg/mL. The natural fibrin clot is strengthened by platelets. In literature, the fibrin matrix is typically prepared at high protein concentration, such as >25 mg/mL F1, to provide sufficient stability and strength^50^. However, our previous research found human cells typically did not grow well in fibrin matrix >9 mg/mL due to the dense fiber structures^50^. Thus, we recommended using <9 mg/mL fibrin for 3D *in vitro* cell culture, cell delivery and wound healing application^50^. However, at such low concentration, it required FXIII-catalyzed covalent crosslinking to form a stable fibrin matrix^50^. To achieve this we expressed a recombinant FXIII^50^ with a specific cross-linking activity of 154 U/mg which was about 6-fold higher than plasma-derived FXIII (**Fig. 1b**).

We then tested if stable fibrin matrix could be formed with low F1 concentration using this recombinant FXIIIa, and how the pFN influenced the gelation kinetics and resultant matrix mechanical properties. We used a commercially available, biotherapeutic grade recombinant thrombin and our rFXIIIa at different concentrations to make fibrin matrices with γγF1 alone, γγ’F1 alone, γγ’F1:pFN complex, or a 1:1 mixture of γγF1 and pFN. All sealants were formulated at an F1 concentration of 2.5 mg/mL with 3.3 mg/mL pFN for both γγ’F1:pFN and analogous γγF1+pFN mixtures. All groups formed stable matrices with higher concentrations of rFXIIIa observed to accelerate gelation and improve the mechanical properties of the fibrin (**Fig. 1c-f**). Thus, this highly active rFXIIIa allowed us to use a low concentration fibrin matrix for our following *in vitro* cell culture and *in vivo* studies.

### A 3D fibronectin nano-array was formed on the γγ’F1:pFN matrix

We used immunostaining and confocal microscopy to study the distribution of pFN and F1 in the fibrin matrices (**Fig. 2**). We found that all formulations produced similar fibrin fiber networks. Most fibers were between 100 and 400 nm in diameter while having a similar degree of branching. Large depots of pFN sporadically occurred within the fibrin matrix made from the γγF1+pFN mixture. The fibrin made from the γγ’F1:pFN complex presented pFN throughout the matrix as a starkly periodic, banded structure. This banded structure appeared superimposed on the continuous γγ’F1 fiber network.

**Fig. 2.**
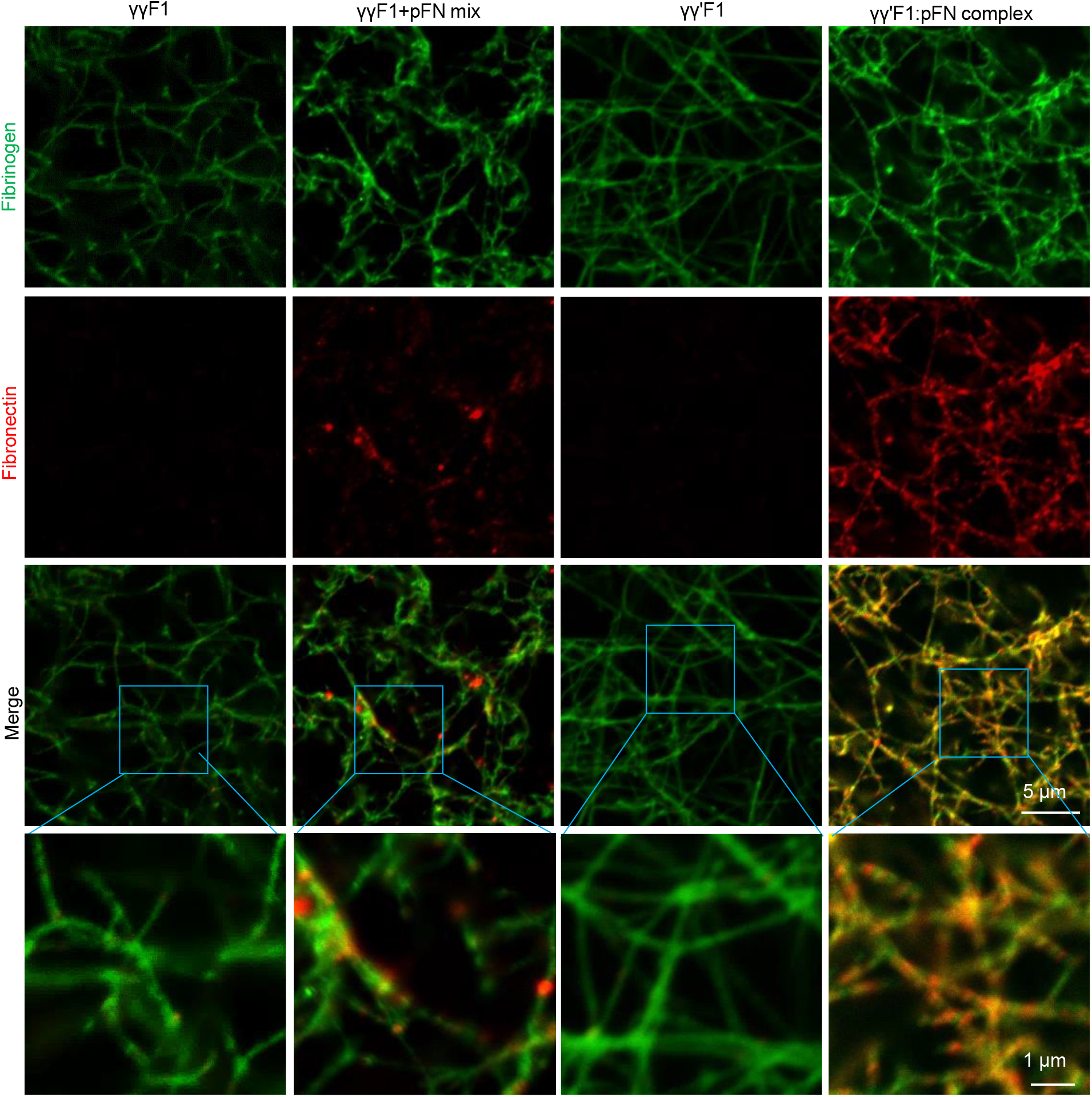
Confocal microscopy fibrin matrices made from γγF1, γγ’F1, 1:1 mixture of γγF1 and pFN, or γγ’F1:pFN complex.

### The 3D fibronectin nano-array enhanced the adhesion, migration, proliferation, and function of healing cells

As previously documented, healing cells interact with the fibrin provisional matrix through binding fibronectins with their integrin receptors^32–35^. We hypothesized that the 3D fibronectin nano-array could potentiate the cell-matrix interaction. We first tested the interactions of fibroblasts with these 4 different fibrin matrices. The adhesion of fibroblasts to fibrin made with γγF1, γγ’F1, 1:1 γγF1+pFN mixture, and γγ’F1:pFN complex were examined through culturing them on top of the matrix. Cells were allowed to attach to the matrix for 3 hours, a time that is sufficient for cellular adhesion. The nuclei of the attached fibroblasts were stained by DAPI (**Fig. 3a**). Adhered cells were also quantified with Alamar blue assay (**Fig. 3d**). The results showed that fibroblasts had similar adhesion to these matrices.

**Fig. 3.**
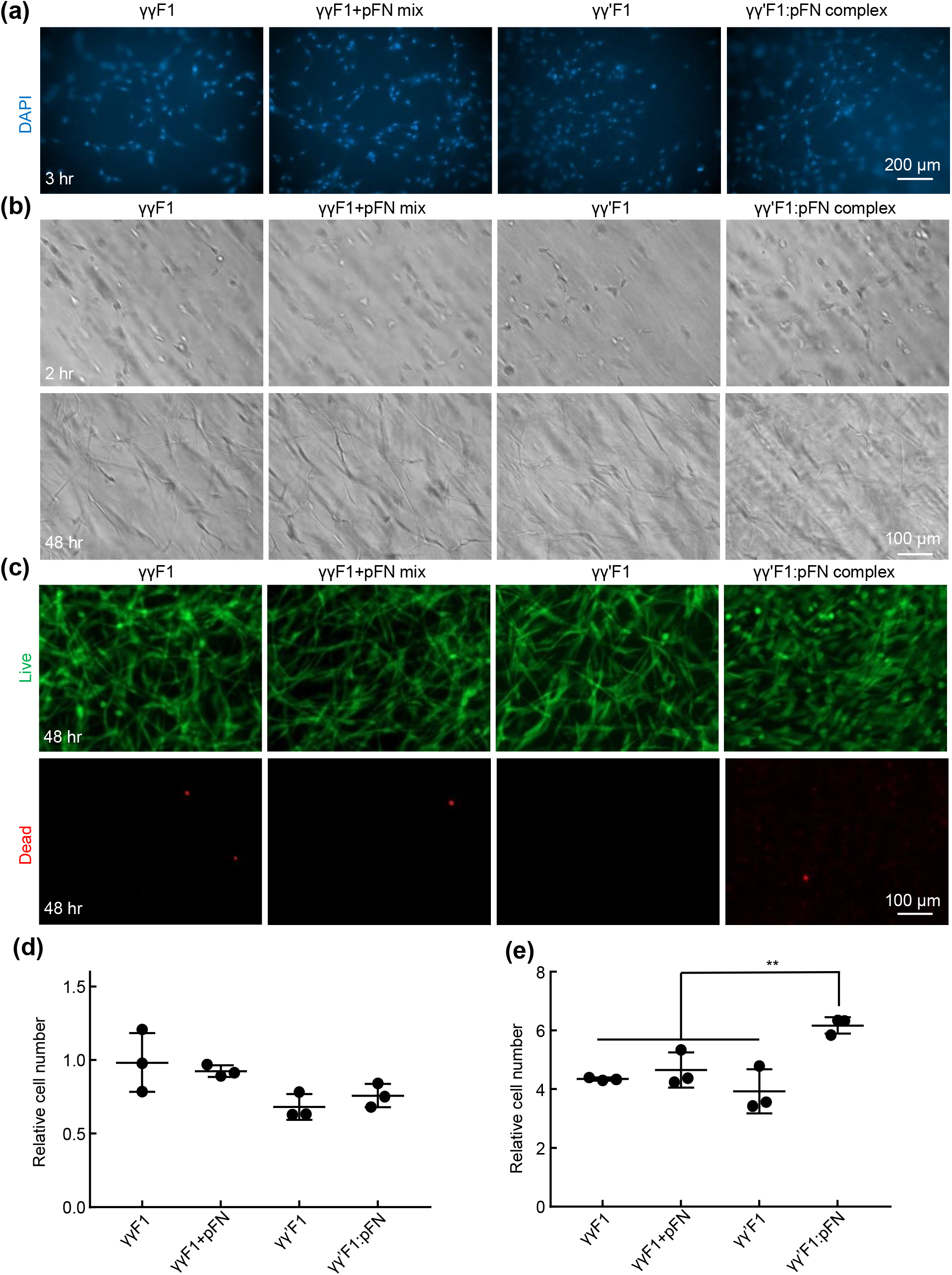
Culturing primary human fibroblasts on top of (a, d) and within 3D fibrin matrices (b, c, e). a) The nuclei of cells attached to fibrin matrices at 3 hr. b) Fibroblasts in 3D fibrin matrices at 2 hr and 48 hr. c) Live and dead cells at 48 hr. d) Cells attached to the fibrin matrices were quantified with alamar blue assay. e) Cells in 100 μl fibrin matrices at 48 hr were quantified with alamar blue assay. Cell numbers are related to the initial seeded cells. **: p<0.01

For efficient wound healing, the provisional matrix also needs to support the survival, proliferation, and morphogenesis of both fibroblasts and endothelial cells (ECs) in addition to good adhesion. Since cells *in vivo* are in the 3D matrix, we also used 3D culture to study these phenomena in the 4 matrices. Fibroblasts were cultured within the fibrin matrix. After 48 hours, fibroblasts exhibited extended morphology, indicating good adhesion in all matrices (**Fig. 3b**). Fibroblasts survived well in all matrices as shown by the live/dead cell staining (**Fig. 3c**). However, the numbers of cells in the γγ’F1:pFN complex matrix was significantly higher (~1.5 fold) than in other matrices (**Fig. 3e**). These results showed the γγ’F1:pFN matrix or the 3D pFN nano-array promoted the proliferation of fibroblasts.

The interactions of HUVECs with these 4 matrices were also investigated. HUVECs adhered to the surfaces of all fibrin matrices after 3 hours. The numbers of HUVECs adhered to the γγ’F1:pFN matrix was 2 fold of these to the γγF1, γγ’F1, and 1:1 γγF1+pFN mixture matrix, indicating HUVECs had stronger adhesion to γγ’F1:pFN complex matrix (**Fig. 4a, and 4d**). After 48 hours of culturing in 3D fibrin matrices, HUVECs in γγF1, γγ’F1, and γγF1 +pFN matrix showed spherical morphology, indicating poor adhesion to the matrices. Some cells showed blebs, a sign of apoptosis (**Fig. 4b**). The existence of apoptosis was confirmed by live/dead cell staining (**Fig. 4c**). HUVECs in γγ’F1:pFN complex matrix showed no apoptosis. Also, they showed a high level of morphological sprouting. Quantification found 400% more cells in the γγ’F1:pFN matrix than other formulations (**Fig. 4e**). These results showed the γγ’F1:pFN complex matrix or the 3D pFN nano-array significantly promoted the adhesion, survival, proliferation, and morphogenesis of HUVECs.

**Fig. 4.**
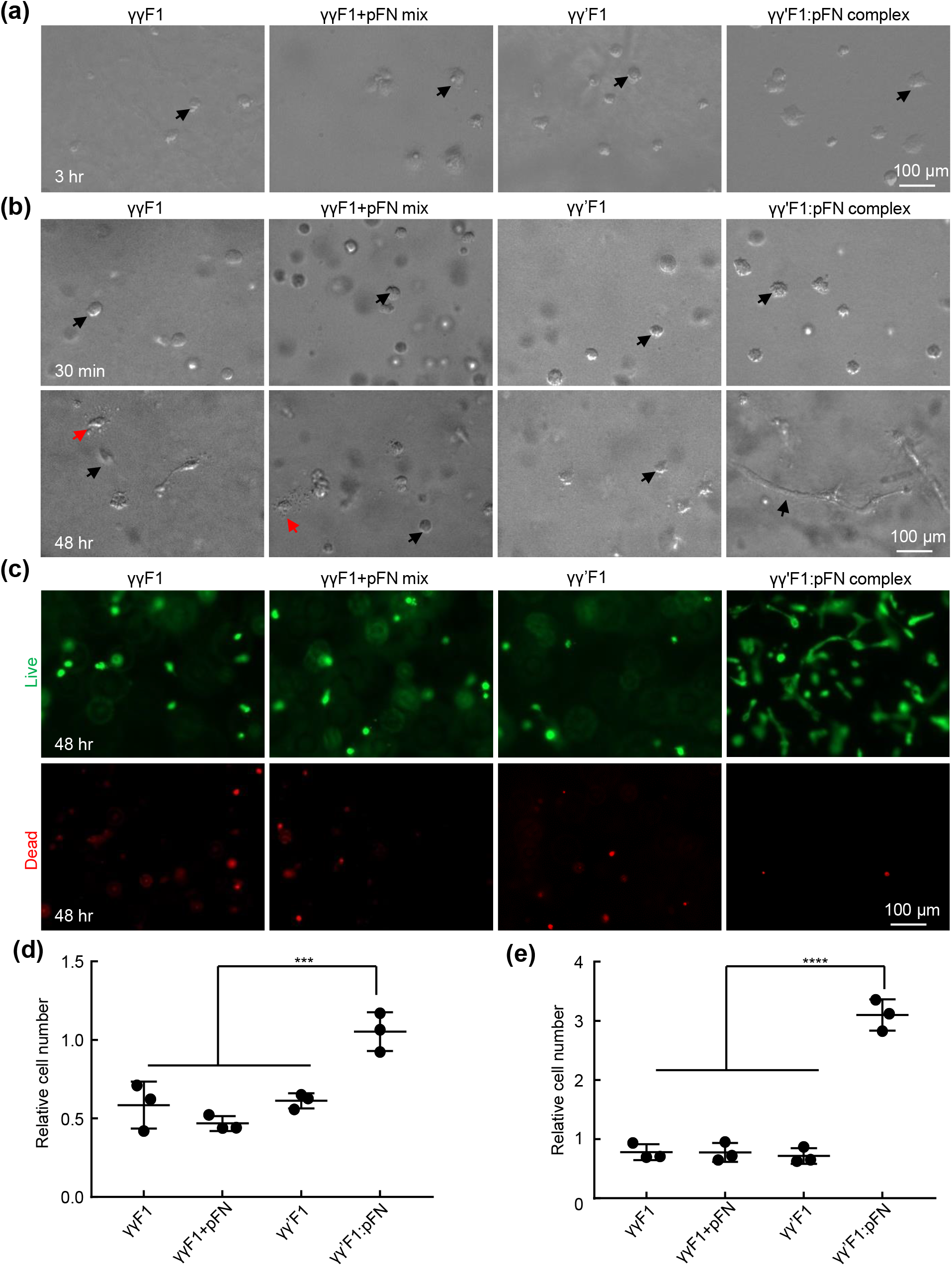
Culturing primary human umbilical vein endothelial cells (HUVECs) on top of (a, d) and within 3D fibrin matrices (b, c, e). a) HUVECs attached to fibrin matrices at 3 hr. b) HUVECs in 3D fibrin matrices at 30 min and 48 hr. Red and black arrows point to apoptotic and live cells respectively. c) Live and dead cells at 48 hr. d) Cells attached to the fibrin matrices were quantified with alamar blue assay. e) Cells in 100 μl fibrin matrices at 48 hr were quantified with alamar blue assay. Cell numbers are related to the initial seeded cells. ***: p<0.001 and ****: p<0.0001.

### The 3D fibronectin nano-array accelerated skin wound healing in healthy mice

Based on the *in vitro* cell culture data, we hypothesized that the novel matrix could enhance the granulation tissue formation and wound closure *in vivo*. We tested this hypothesis in a full-thickness model in healthy mice. Wounds were treated with γγ’F1:pFN complex, or γγF1 and pFN mixture. The macroscopic evaluation showed the γγ’F1:pFN complex significantly speeded up wound closure (**Fig. 5a**). Indeed, 50% and 81% of wounds treated with the γγ’F1:pFN complex achieved full re-epithelization on days 4 and 6 respectively. On the other hand, 22% and 64% of wounds treated with the mixture achieved full re-epithelization on days 4 and 6 respectively, and 0% and 30% of the control wounds (without treatment) achieved full re-epithelization on day 4 and 6 respectively (**Fig. 5c**). The microscopic evaluation confirmed these observations. The migration of epidermal cells was significantly increased in the γγ’F1:pFN complex treated group just (i.e. the leading edge) 2 days post-treatment (**Fig. 6a**) and showed a complete epithelization over the wound bed on day 6 (**Fig. 6b**). Histology also showed dense granulation tissue at the wound center only in the γγ’F1:pFN complex treated group (**Fig. 6b**). These data supported that the novel fibrin sealant/matrix or the 3D pFN nano-array promoted epidermal cell migration, resulting in faster re-epithelization *in vivo*.

**Fig. 5.**
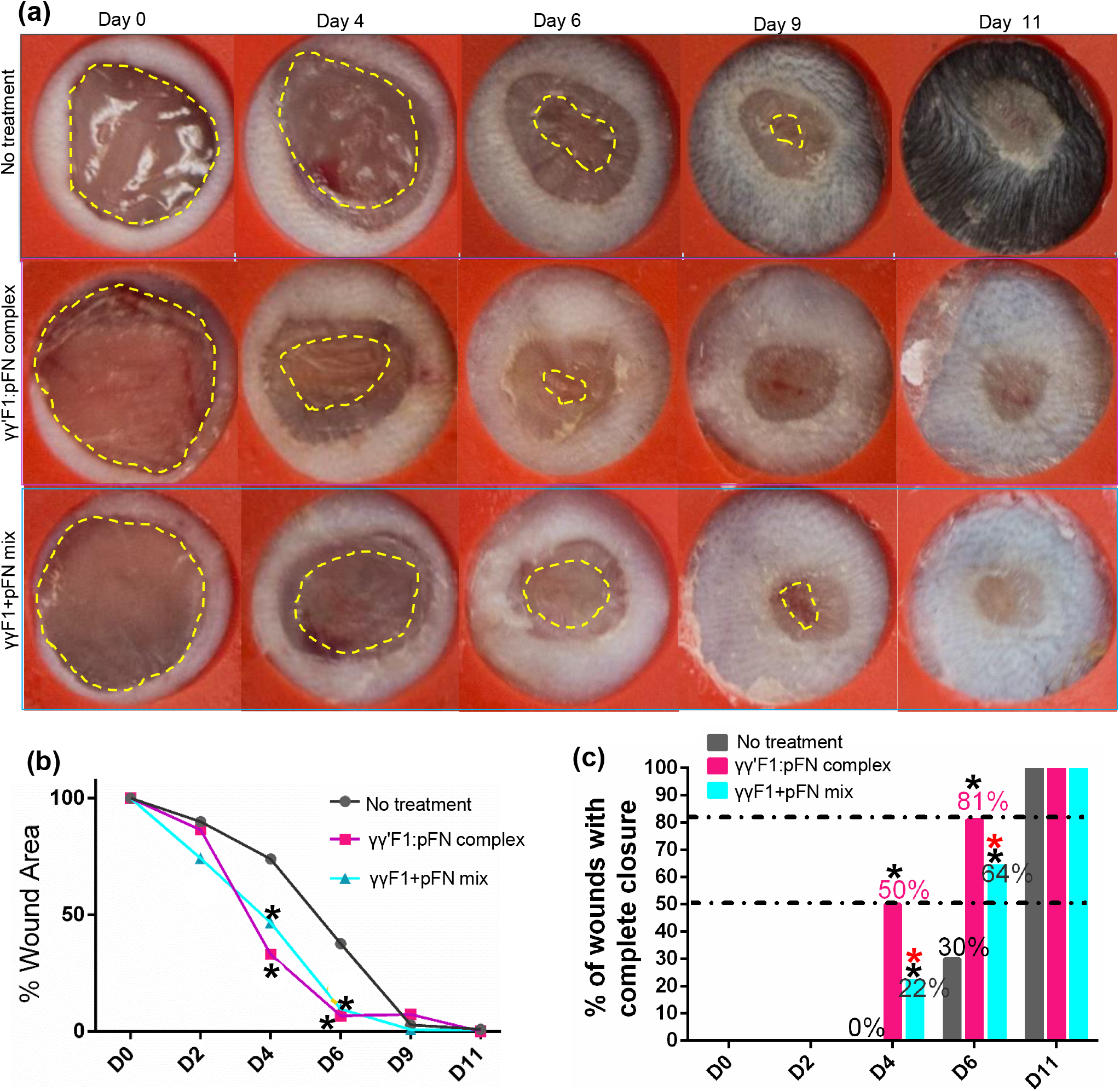
(a) Macroscopic wound morphology on different days. Yellow dash line highlights the leading edge of the new epithelium. (b) The percentage of wound area, and (c) the percentage of mice achieved 100% re-epithelization on different days. *: p<0.05; *:compared to no treatment group; *: compared to complex.

**Fig. 6.**
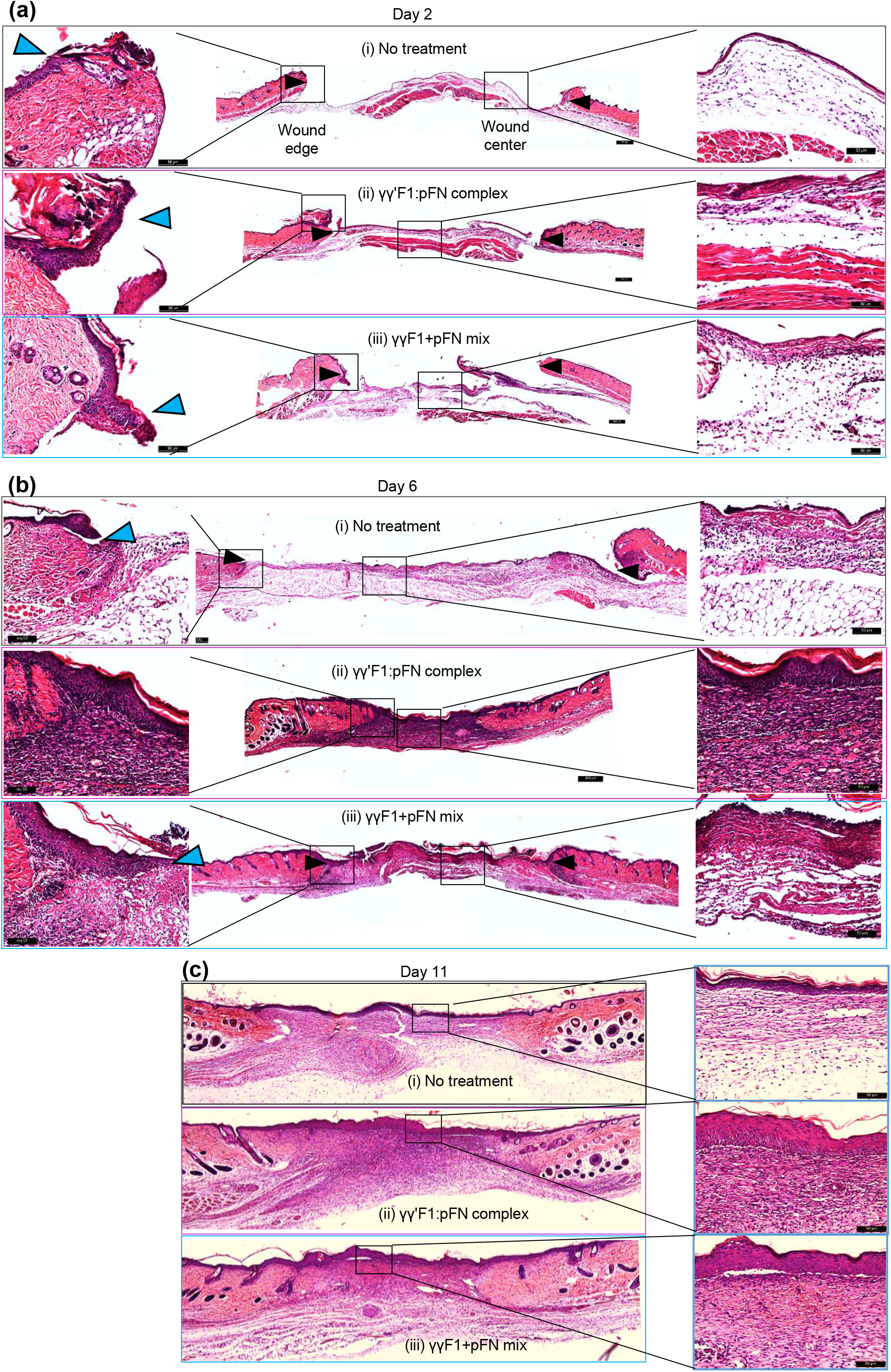
H&E staining of days 2, 6 and 11 wounds.

We also evaluated the extracellular matrix formation via histocolorimetric analysis of collagen content (**Fig. 7**). Total collagen deposition was identified by Masson’s trichrome staining on days 6 and 11 post-wound, period when the highest production of collagen is reached^56^. Animals treated with γγ’F1:pFN complex or γγF1+pFN mixture treatments showed no differences in total collagen deposition. However, the quality of collagen was different. Staining showed dense and aligned reticular collagen fibers in the γγ’F1:pFN complex group on day 6, suggesting high-quality granulation tissue (**Fig. 7a**). The other treatment groups had a distinctly more sparse and randomly oriented collagen layer showing less mature granulation tissue.

**Fig. 7.**
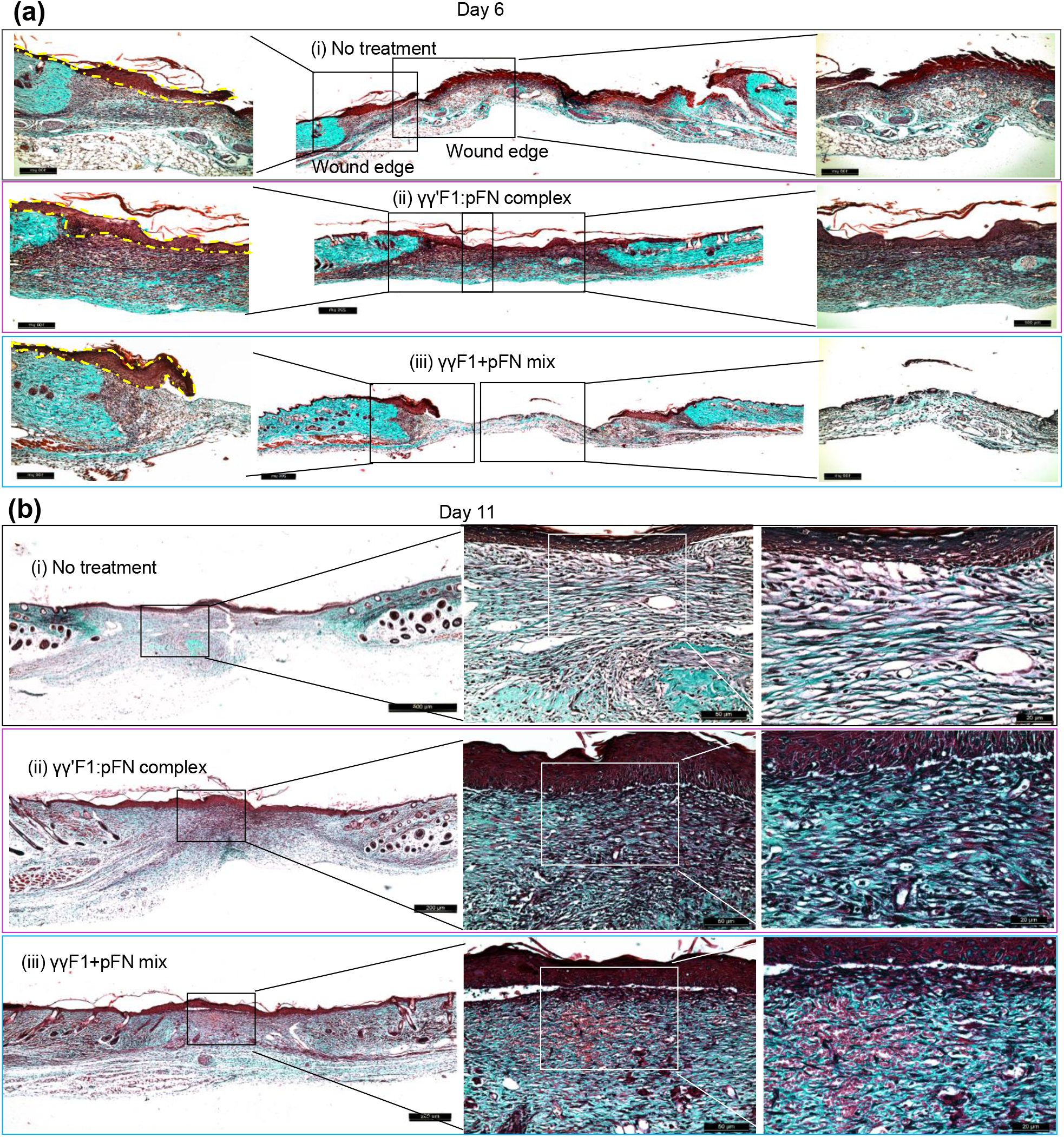
Masson’s trichrome staining was done to evaluate the total collagen deposition. Collagen (blue) on the wound bed was much less than the surrounding normal skin.

We evaluated the collagen subtype progression through Picrosirius Red Staining, which stains collagen III in green and collagen I in red (**Fig. 8**). In physiological wound healing, granulation tissue mainly consists of collagen III in the proliferative phase that gradually changes to mesh-like collagen I in the remodeling phase. Our data of control wounds agreed with this (**Fig. 8**). However, wounds treated with the γγ’F1:pFN complex on day 6 showed higher collagen I content. Even more, γγ’F1:pFN complex on day 6 was similar to day 11 of control group wounds (**Fig. 8a-c**). On day 11, the γγ’F1:pFN complex group had the mesh-like collagen organization (**Fig. 8b**). These data showed the γγ’F1:pFN complex shifted the granulation formation, collagen remodeling about 5 days earlier, and improved the quality of the new tissue.

**Fig. 8.**
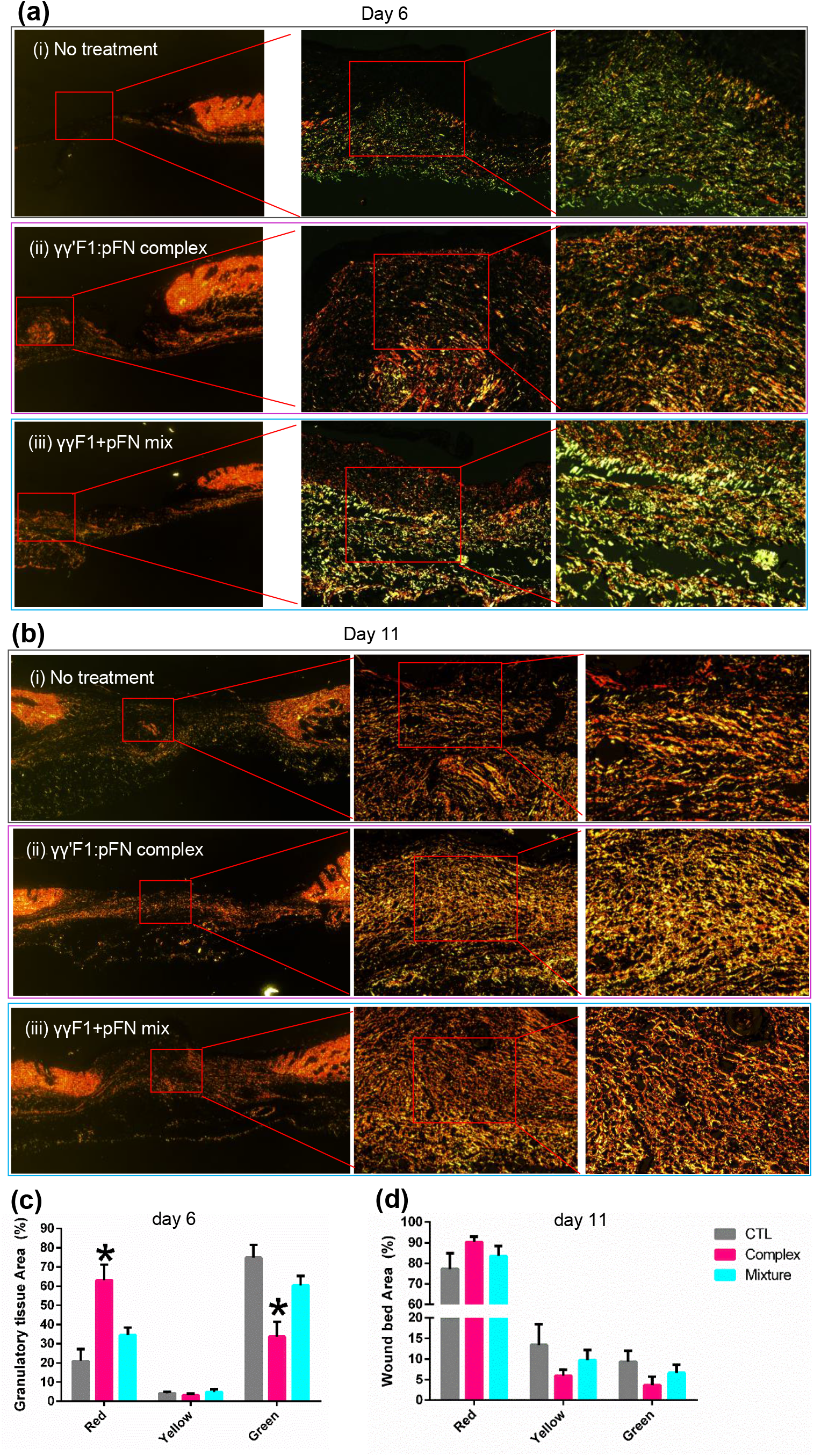
Picrosirius red staining on collagen I (red) and III (green) on day 6 (a, c) and 11 (b, d). *:p<0.05

We assessed the expression of 20 signature genes associated with the transitioning between sequences of inflammation, cell migration, and proliferation^57^ that could corroborate macroscopic changes in dermal tissue structure. Several statistically significant changes in gene expression related to the sequence of these processes were observed among the broad panel of assays presented here (**Fig. 9a**). For signals associated primarily with an active inflammatory phase, we observed by day 2 the γγ’F1:pFN fibrin treatment group had decreased expression of inflammatory cytokines IL-β1 and IL-10 relative to the other treatment groups(**Fig. 9a**). Furthermore, by day 6 both relative levels of IL-β1 and IL-6 had receded to very low relative levels as had the macrophage marker F4/80 and monocyte chemoattractant MCP-1. These already diminished signal profiles related to the inflammatory phase indicates that the inflammatory phase was resolved very early in the healing process.

**Fig. 9.**
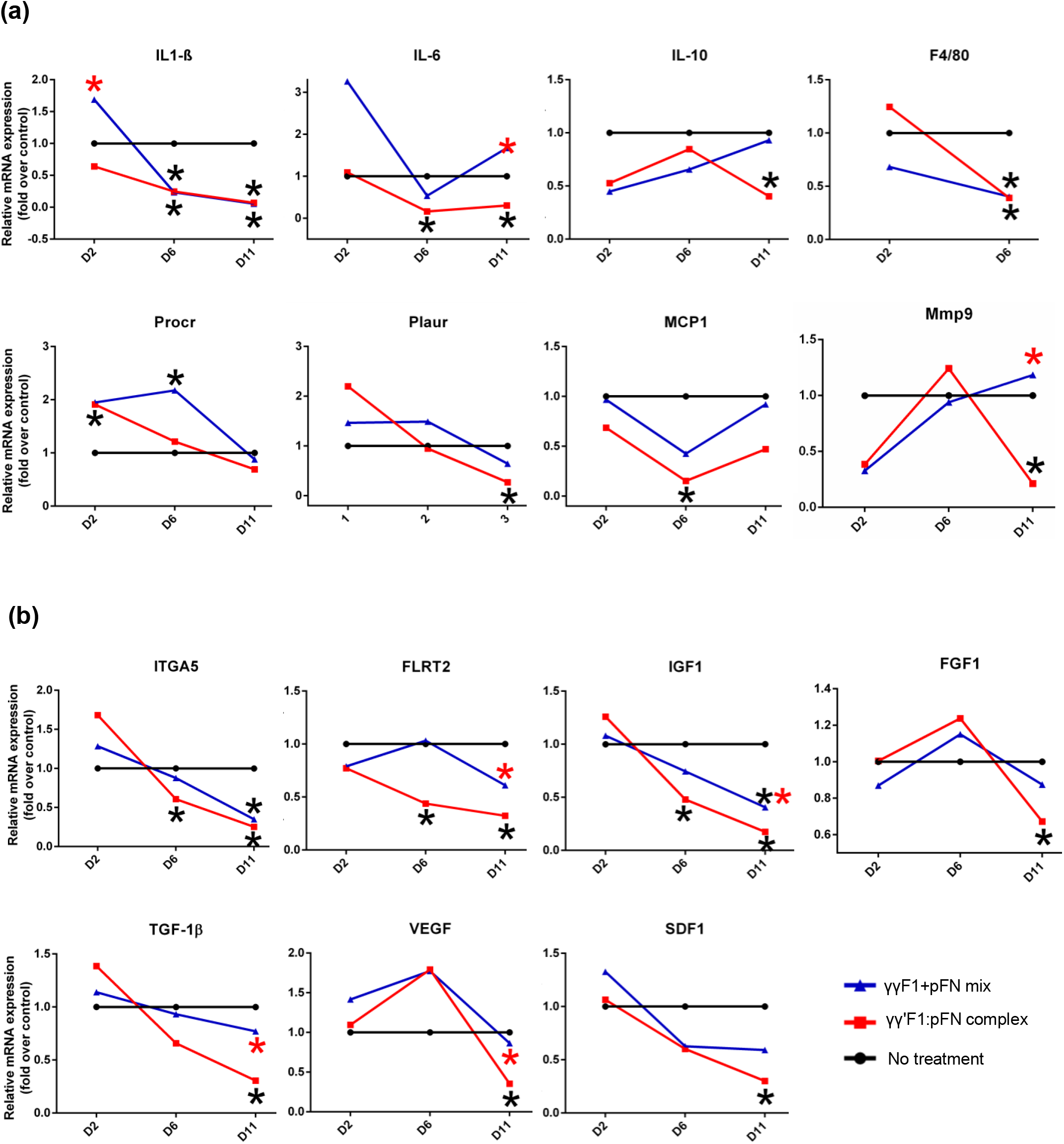
The relative expression of genes related to inflammation (a), and cell migration and proliferation (b). RNA level was normalized to the untreated group. *:p<0.05, *:compared to no treatment group; *: compared to complex.

The transition from inflammation to signaling needed for keratinocyte migration also occurred early. We found that PROCR, a gene highly expressed by migrating keratinocytes^58^, was increased by the complex and Mixture treatments on day 6 (**Fig. 9a**). Moreover, the complex group showed an early expression of PROCR on day 2. The Fibronectin Leucine Rich Transmembrane Protein 2 (FLRT2), a cell-cell adhesion marker for keratinocytes, is highly expressed in the leading edge during wound healing^57^. FLRT2 is not expressed in normal skin epidermis but is expressed during the granulation tissue formation. Our result showed that the FLRT2 expression was decreased on days 6 and 11 in the γγ’F1:pFN complex group, supporting that the cell migration was completed before day 6 in the complex group (**Fig. 9b**). Keratinocytes bind to the fibrin matrix using alpha 5 integrins (ITGA5), in order to migrate. We found ITGA5 expression increased on day 2 and significantly decreased on days 6 and 11 for the complex group, supporting that cell migration was completed before day 6 (**Fig. 9b**). These data agree with the acceleration of tissue structure observed such as the complete re-epithelization and granulation tissue structure found on day 6 for the γγ’F1:pFN treatment group.

Growth factor signaling is critical for granulation formation. We found the γγ’F1:pFN treatment decreased expression of TGF-β1, FGF1, SDF1, VEFG-α and IGF1 genes on day 11, supporting that cell proliferation, granulation tissue formation, and re-epithelization were completed before day 11 in the γγ’F1:pFN complex group (**Fig. 9a**). Even more, IGF1 early declined on day 6, showing the end of the proliferative phase. Taking all together, animals treated with the γγ’F1:pFN complex showed a noninflammatory, non-proliferative, and non-migrative profile on day 11, suggesting an early resolution of the wound healing process. In short, our data showed the γγ’F1:pFN complex or the 3D pFN nano-array greatly accelerated skin wound healing in healthy mice.

## Discussion

The topical application of the fibrin matrix represents a promising approach to enhance the healing of skin wounds. The natural provisional matrix contains a small number of fibronectins^1,59–61^. Previous reports have shown that raising the fibronectin content in the matrix promotes wound healing. For instance, human fetus^62–64^ and invertebrate animals newts^65^ heal wounds much faster because their provision matrices contain much more fibronectins that allow cells to migrate and proliferate faster. The topical application of pure soluble fibronectins enhances normal and diabetic wound healing in animals^62,66–70^. These soluble fibronectins aggregate or attach to surrounding tissue, but do not form a 3D matrix in the wound bed. A recent study shows that if processed into a 3D nanofiber scaffold, fibronectins can heal normal skin wound much faster^62^. However, processing fibronectin nanofibers at a large scale is extremely challenging, limiting their clinical applications.

Both receptor occupancy and clustering are required for integrin signaling^71–73^. Most recent research shows that one way to boost integrin signaling is to present fibronectins as nanoclusters that can cluster the integrin receptors. For example, when cells are cultured on a uniformly fibronectin-coated substrate, high fibronectin density is required to trigger the integrin signaling. On the other hand, if fibronectins are patterned into nanoclusters, they trigger a much higher integrin clustering, signaling, and cell function^73–79^ However, nano-patterning fibronectin has only be achieved on a 2D substrate to date. How to pattern fibronectins as nanoclusters in a 3D matrix, especially on the fibrin nanofibers, and how the 3D nanoscale presentation influences the cell-matrix interaction and cell signaling are still unknown.

In our previous paper^8^, we reported a bioprocess to separate γγF1 and γγ’F1 from plasma and we found that γγ’F1 and pFN formed a tight complex. The γγ’F1 and pFN occur at nearly a 1:1 molar ratio in normal human plasma^9,10,80,81^, thus the γγ’F1:pFN complex we isolated may already exist in human plasma (Fig. 1). The γγ’F1:pFN complex formed a matrix with fibronectin nanobands on the fibrin nanofibers in the presence of highly active thrombin and rFXIIIa (Fig. 2). We showed that the fiber diameters and branch points are markedly similar for all of the fibrins studied. Thus, the fibronectin nanobands likely arose from interactions of pFN with the γ’ tails that are uniquely displayed by the γγ’F1 fibrils. Future research should make clear the dynamics of forming these nanobands, as well as how the sealant composition affects the matrix structure and properties. This report successfully reproduced all our previous findings (Fig. 1, 2), showing the robustness of the separation process, the γγ’F1:pFN complex, and the novel nanostructures.

We found that the γγ’F1:pFN matrix potentiated the adhesion of key cells that participate in the wound healing process including fibroblasts (Fig. 3) and endothelial cells (Fig. 4) *in vitro*. This enhanced interaction led to boosted cell survival, proliferation, migration, and morphogenesis in 3D culture. This was particularly clear for endothelial cells. Because of the similarities in core fiber matrix structure among all fibrins studied here, the enhanced interaction of ECs with γγ’F1:pFN fibrin was likely due to the pFN nanobands. As said, angiogenesis is the rate-limiting step of granulation tissue formation, and granulation tissue formation is the rate-limiting step of wound healing. Thus the capability to stimulate endothelial cells has a significant impact on wound healing. The plasma-derived fibrin has a 10:1 ratio of F1 to pFN^7^. This fibrin has been previously shown to support the colonization of fibroblasts^82^, but not EC. FGF-2 and or VEGF supplementation has been reported as the primary catalyst of EC colonization in plasma-derived fibrin^83–85^. In contrast, we observed an accelerated EC colonization in the γγ’F1:pFN fibrin without using these growth factors. Future research should systematically make clear the dynamics of cell-matrix interaction and integrin clustering, and how the interaction alters the global gene expression and cell phenotypes. It also is very valuable to systematically study how the sealant composition and matrix structure influence the matrix interaction with the major cell types of the wound healing process.

Our *in vivo* studies agreed well with the *in vitro* cell culture results. The novel nanostructure significantly accelerated the re-epithelization (Fig. 5, 6), granulation tissue formation, and modeling (Fig. 7, 8). The whole wound healing process was speeded up about 5 days. Additionally, the quality of the granulation tissue was significantly improved. This conclusion was supported by the macroscopic morphology (Fig. 5) and microscopic structure analysis (Fig. 6, 7, 8) as well as molecule profiles (Fig. 9). Here we have shown that animals treated with γγ’F1:pFN complex accelerated cell migration. The previous report showed that cells at the leading edge in direct contact with the fibrin matrix do not proliferate actively but migrate to the center of the wound^57,86^. Animal treated with the γγ’F1:pFN complex modulated two key membrane proteins that guide cell migration and cell adhesion. It will be valuable to systematically study the cellular profiles at different time points along the healing process in the future.

As said, large acute skin wounds after surgery, trauma and burn heal slowly (e.g. take >1 month) and result in scarred skin that has an abnormal appearance, structure, and function. We believe our novel matrix has a high potential to promote the healing of these wounds. Since all materials used in our matrix are human proteins, their safety is expected. Thus our matrix can be quickly translated to bedside once the efficacy and underlying mechanisms are made clear. Since our matrix is fibrin-based, it can be used for both hemostasis and enhancing wound healing. In addition to the acute wounds, our matrix may be used to treat the chronic wound as well. Skin wounds in the aged population, people with diabetes, obesity, and vascular diseases present serious difficulties to heal^87,88^. Current treatments have limited success^87–90^. It will be very valuable to study treating chronic wounds with our novel sealant in the future.

In conclusion, we confirmed that the 10% γγ’F1 in plasma could be separated from the 90% γγF1 using our previously reported method^8^, as well as γγ’F1 formed a complex with pFN. We also confirmed that the γγ’F1:pFN complex formed a fibronectin-nanobands-on-fibrin-nanofiber structure. We found that the 3D nano-scale presentation of the FN could significantly potentiate the cell-matrix interaction *in vitro* and accelerate the granulation formation, re-epithelization in a mice skin wound.

## Acknowledgments

This research was supported by the Nebraska Collaboration Initiative; the Nebraska DHHS Stem Cell Grant; the US Army Medical Research Acquisition Activity (W81XWH-05-1-0527); and the Department of Defense, USA (W81XWH-BAA-11-1). This study was financed in part by the Coordenação de Aperfeiçoamento de Pessoal de Nível Superior – Brasil (CAPES) – Finance Code 88882.434714/2019-01. Confocal microscopy imaging was done in the Morrison Microscopy Core Research Facility at the University of Nebraska, Lincoln.

## Competing interests

N/A

## Notes

### Competing Interest Statement

The authors have declared no competing interest.

